# Soil and atmospheric drought trigger early leaf senescence and increase subsequent canopy mortality risk in temperate forests

**DOI:** 10.64898/2025.12.18.694887

**Authors:** Pia Labenski, Allan Buras, Rüdiger Grote, Martin Thurner, Nadine K. Ruehr

## Abstract

Early leaf senescence – an advanced decline in leaf functionality before the typical autumn period – has emerged as recurring drought response in temperate forests. Yet its large-scale patterns, drivers, and impacts on tree vitality remain poorly understood. Using six years (2018–2023) of Sentinel-2 time series across Germany, we estimated the onset of leaf senescence in European beech (*Fagus sylvatica*) and oak (*Quercus robur, Q. petraea*) forests. Early leaf senescence was most widespread during exceptional drought years (2018, 2019, and 2022) and showed pronounced spatial variability. Sustained high atmospheric demand (VPD) and critically low soil water potential (ψ_soil_) in summer were key drivers identified by logistic regression analysis (F1-score = 0.57), with beech responding more strongly to soil drought and oak to atmospheric drought. Thresholds triggering early leaf senescence (probability >50%) were six weeks of daily maximum VPD ≥1.9 kPa in beech and ≥2.1 kPa in oak, or two weeks of root zone ψ_soil_ ≤-0.8 MPa and ≤-0.9 MPa, respectively. Elevated spring VPD further amplified early senescence risk in both species, indicating seasonal legacy effects that may reduce hydraulic function or leaf longevity. Forests with very early and repeated early leaf senescence exhibited elevated canopy mortality in subsequent years, highlighting them as potential warning signals of drought-induced forest decline. K-means clustering of forest stands based on senescence patterns, mortality, and drought sensitivity classified three drought response types – resistant, sensitive, and vulnerable – shaped mainly by local soil water availability and stand structure. Our findings demonstrate that early leaf senescence reflects cumulative drought stress and indicates increased mortality risk, rather than a stress-avoidance strategy. We propose it as a mechanistic indicator that can be incorporated into ecosystem models to improve predictions of forest vulnerability and carbon dynamics under climate extremes, and guide forest management to anticipate climate-induced forest decline.

## 1. Introduction

Between 2018 and 2022, European forests experienced an unprecedented series of hot droughts (Allen et al., 2015), marked by low precipitation, above-average air temperatures, and high evaporative demand in multiple years, leading to persistent soil moisture deficits (Bevacqua et al., 2024; van der Wiel et al., 2023). In response to this intensifying compound stress, declines in forest health were observed across the continent, with particularly high severity in Central Europe (George et al., 2022; Knutzen et al., 2025; Schuldt et al., 2020). One of the earliest visible symptoms was the widespread occurrence of premature leaf coloration and shedding in deciduous trees during the summer of 2018, in some areas starting already in late July (Buras et al., 2020; Obladen et al., 2021; Schuldt et al., 2020). Large-scale remote sensing analyses estimated that this phenomenon, referred to as early leaf senescence, affected 11% of Central European forests that year (Brun et al., 2020) and similar events recurred in subsequent summers (Descals et al., 2023), although their extent has yet to be quantified. Such shifts in leaf phenology can profoundly impact ecosystem processes by altering the timing and magnitude of carbon, water, and nutrient fluxes (Richardson et al., 2013). Yet, the specific stress conditions that trigger early senescence and its long-term consequences for tree health remain insufficiently understood.

The response of leaf senescence to increasingly warm and dry conditions is highly variable. While moderate warming often delays senescence in temperate species (Delpierre et al., 2009; Grossiord et al., 2022; Vitasse et al., 2011; Xie et al., 2018), evidence suggests that seasonal effects differ.

Increased spring temperatures can advance senescence, potentially due to shortened leaf lifespan following early leaf-out, enhanced productivity leading to sink limitation, or cumulative drought stress later in the year (Fu et al., 2014; Keenan & Richardson, 2015), whereas warmer summer and autumn conditions tend to delay senescence (Fu et al., 2018; Zohner et al., 2023). Under extreme conditions, however, marked by strong atmospheric and soil moisture deficits, unusually early leaf senescence has been repeatedly observed in temperate forests (Bigler & Vitasse, 2021; Bréda et al., 2006; Schuldt et al., 2020; Walthert et al., 2021). High temperatures, declining precipitation amount and frequency, and drought severity have been identified as common drivers of earlier senescence in large-scale studies (Brun et al., 2020; Descals et al., 2023; Ma et al., 2024; X. Zhang et al., 2025), but responses are not fully consistent across all regions or species. For instance, during the extreme drought of 2003, both earlier and delayed senescence were observed in deciduous forests in Central Europe (Bréda et al., 2006; Leuzinger et al., 2005), likely reflecting species-specific strategies, local site conditions, and variation in inter- and intraspecific physiological thresholds (e.g., Zuccarini et al., 2023). Forest composition and structure, soil properties, and terrain further modulate senescence responses at the regional scale (Brun et al., 2020; Meier et al., 2021).

Different physiological mechanisms can induce early leaf senescence under drought, spanning from adaptive responses to stress-induced damage. As an adaptive, stress-avoidance strategy, early leaf senescence can limit water loss and protect the hydraulic system from embolism, albeit at the cost of reduced carbon assimilation (Leuschner, 2020; Nadal-Sala et al., 2021; Ruehr et al., 2016; Wolfe et al., 2016). Conversely, early senescence may also reflect cumulative damage, for instance caused by fast dehydration processes at the leaf level resulting in xylem embolism, or direct heat-induced tissue damage (Scafaro et al., 2021). In temperate forests, drought-induced leaf shedding is well documented in European beech (*Fagus sylvatica*) (Bréda et al., 2006; Leuschner, 2020), whereas oak responses are more variable. Experimental studies on oak saplings showed both advanced and delayed senescence in response to drought, depending on drought intensity, timing, and the presence of re-watering (Günthardt-Goerg et al., 2013; Spieß et al., 2012; Vander Mijnsbrugge et al., 2016). In mature oaks, an advanced loss of canopy greenness under drought has been observed in *Quercus robur* (Mariën et al., 2021; Zuccarini et al., 2023), and strong defoliation was reported in sub-mediterranean *Quercus pubescens* under extreme drought (Pollastrini et al., 2019). For European beech, the onset of drought defoliation in mature trees has been linked to critical thresholds of predawn leaf and soil water potential across forest plots spanning a range of water availability conditions in Switzerland (Walthert et al., 2021). However, such stress thresholds have not yet been identified at larger spatial scales and are currently lacking for temperate oaks.

In relation to the proposed mechanisms of early leaf senescence above, there is some uncertainty about whether early leaf senescence should be interpreted as a sign of drought resistance or as indicator of increased drought vulnerability and precursor for long-term decline. For instance, beech forests in France that exhibited early senescence during the 2003 summer drought did not show crown deterioration in the following year (Bréda et al., 2006). However, more recent evidence from European beech suggests that early senescence often indicates drought-induced damage, as increased crown dieback and lagged tree mortality have been documented in early senescing beech trees in the years following the 2018 summer drought (Frei et al., 2022; Walthert et al., 2021; Wohlgemuth et al., 2020). This has been linked to persistent drought-induced xylem embolism, limiting water supply and reducing leaf area as well as overall canopy health in the following year (Arend et al., 2022). If hydraulic function remains impaired or defoliated trees cannot maintain a positive carbon balance, a steady decline of tree vigor can be expected particularly under recurrent droughts (Hájíčková et al., 2024; Salmon et al., 2015). Thus, early leaf senescence may serve as an early-warning signal of tree decline and increased mortality risk in subsequent years (Frei et al., 2022; Hunziker et al., 2024).

Given the high spatial and temporal variability in forest drought responses, remote sensing offers an effective means to monitor early senescence at large scales. High-resolution Sentinel-2 time series have proven useful in capturing patterns of early leaf senescence across Central Europe (Brun et al., 2020; Descals et al., 2023), but these studies relied on the normalized difference vegetation index (NDVI), which saturates in dense canopies (LAI >2-3 m² m^-2^; Gao et al., 2023) and may thus limit sensitivity to subtle changes in crown condition. Sentinel-2 red-edge bands, by contrast, remain sensitive to both chlorophyll content and LAI even in dense canopies, potentially making them more effective for detecting early senescence (Mariën et al., 2021; Zarco-Tejada et al., 2018). Notably, satellite-based senescence onset has rarely been validated against ground observations (but see Liu et al., 2024), as these typically capture later stages of senescence (e.g., ≥50% leaf coloration; Tian et al., 2021; Y. Zhang et al., 2024). Additionally, species-specific patterns of early senescence have not yet been quantified at large spatial scales, as wall-to-wall high-resolution tree species information became available only recently (e.g., Blickensdörfer et al., 2024). Finally, large-scale detections of early leaf senescence have mostly been linked to general climate variables such as temperature and precipitation, while more physiologically meaningful indicators of plant drought stress, such as vapor pressure deficit (VPD) and particularly soil water potential, remain underrepresented.

Here, we derive spatial patterns of leaf senescence onset in beech- and oak-dominated forests of Germany during 2018–2023 using time-series of Sentinel-2 red-edge indices, and compare them to ground-based observations from the permanent intensive forest monitoring plots (Level II of ICP Forests, https://www.icp-forests.net/). We quantify the extent of forest areas with early leaf senescence and link these patterns to atmospheric and soil drought indicators (VPD and soil water potential), additional site variables, and tree mortality maps to address the following research questions:

Q1) What is the extent and spatial distribution of early leaf senescence in German beech and oak forests during 2018–2023?

Q2) Which cumulative drought conditions trigger early senescence, and how do atmospheric versus soil drivers differ in strength and timing between species?

Q3) Is early leaf senescence associated with increased canopy mortality in subsequent years, and thus an early indicator of long-term tree decline?

Q4) Can distinct patterns of drought sensitivity in senescence timing be identified, and which environmental or site-specific factors explain them?

## 2. Methods

### 2.1 Leaf senescence onset from satellite data

#### 2.1.1 Sentinel-2 time series processing

We used Level 2 Analysis Ready Data (ARD) from Sentinel-2 at 10 m resolution covering Germany for the years 2018–2023, available via the EO-Lab platform (https://eo-lab.org/). The ARD were generated using the FORCE software (Frantz, 2019), and included cloud and cloud shadow masking, atmospheric, topographic, and additional radiometric corrections, and finally data cubing. For further analysis, we included only forest areas dominated by European beech or oak species (*Quercus robur* and *Quercus petraea,* available only in a combined *Quercus* class) according to a national 10 m tree species map (Blickensdörfer et al., 2024), and with >70% canopy cover according to the Copernicus tree cover density product (Copernicus, 2024), to minimize the influence of understory vegetation on the spectral signal.

Using the FORCE time series analysis module (Frantz, 2020b), we computed four vegetation indices (VIs) – three red-edge-based VIs and the NDVI (Table S1 in the supplementary information, SI) – as candidate VIs for estimating senescence onset (SO). The VI time series were quality-screened using Quality Assurance Information (QAI) flags, and remaining outliers were removed via residual-based filtering with linear interpolation of observation triplets (Frantz, 2020a). The filtered time series were interpolated using a Radial Basis Function (RBF) filter with three different kernel widths to generate weighted moving window averages based on local data availability (Schwieder et al., 2016), resulting in a daily time series. Residual noise was further reduced by fitting piecewise cubic splines that capture seasonal dynamics.

SO was estimated from the smoothed VI time series as the day of year (DoY) when the signal dropped below different candidate seasonal amplitude thresholds (90%, 85%, 80%, 75%, 70%) after the summer maximum. Forest areas affected by disturbance events (fires, windthrows, bark beetle outbreaks, salvage logging) were excluded for the affected year using the European Forest Disturbance Atlas (Viana-Soto & Senf, 2025).

#### 2.1.2 Comparison with ground observations

We compared the remotely sensed SO estimates to phenological observations from ICP Forests Level II plots (Fleck et al., 2016). For this, we extracted the mean SO for the pixels covering beech- and oak-dominated ICP Forests plots (assuming a size of 0.25 ha, the minimum requirement for Level II plots) and correlated these estimates with the field-recorded dates for the onset of coloration and litterfall. These corresponded to the first observation of ‘infrequent or slight’ (1-33% of the canopy affected) coloration or litterfall. We selected the VI and amplitude threshold with the lowest RMSE between the remotely sensed SO and the field-recorded dates. SO maps derived from this VI and amplitude threshold were used in all subsequent analyses.

#### 2.1.3 Early senescence definition and map preparation

We defined early senescence onset (SO_early_) as occurring before DoY 244 (1 September in normal years, 31 August in leap years), consistent with previous studies (e.g., Descals et al., 2023), and classified later dates as normal senescence onset (SO_normal_). We also explored an anomaly-based definition of early leaf senescence similar to Bigler & Vitasse (2021), comparing each pixel’s SO in a given year to its multi-year mean. However, due to missing SO estimates for some pixels in certain years – caused by data gaps exceeding the interpolation window – and the overall short time series, we opted for the fixed-threshold approach.

SO maps at 10 m resolution were used to quantify the forest area affected by early senescence for each species and year (research question Q1). For subsequent analyses involving climate and soil variables (sections 2.2 and 2.5), we aggregated the 10 m SO maps to 1 km resolution to match the spatial resolution of these covariates. This was done by averaging the SO values of beech- or oak-dominated forest pixels within each 1×1 km grid cell. To ensure representativeness, we only computed averages for cells where the respective forest type (beech- or oak-dominated) covered ≥10% of the area, thereby reducing the influence of isolated and potentially noisy pixels. SO_early_ at the 1 km scale was defined based on the aggregated mean DoY before 244.

### 2.2 Atmospheric and soil drought data

We calculated hourly vapor pressure deficit (VPD) for April–August from 2018–2023 using the German Weather Service’s gridded HOSTRADA dataset, which provides near-surface air and dew point temperatures at 1 km resolution based on interpolated station data (DWD, 2024). From these, we extracted daily maximum VPD values (VPD_max_), which typically occur in the early afternoon and are more indicative of drought stress experienced by plants than daily means.

Soil water potential (ψ_soil_) under beech and oak forests for 2018–2023 was also obtained from the German Weather Service as 8-day averages at 1 km resolution (P. Schmidt-Walter, personal communication, April 2025). These data were generated with the hydrological model LWF-Brook90 (Hammel & Kennel, 2001; Schmidt-Walter et al., 2020). LWF-Brook90 estimates soil water potential and moisture dynamics within the species-specific rooting (1.4 m for beech, 1.8 m for oak) using parameters based on Schmidt-Walter et al. (2019) and Weis et al. (2023).

### 2.3 Logistic regression modelling

We used logistic regression to assess the influence of atmospheric and soil drought stress on senescence onset (Q2), treating the outcome as binary variable: 0 for SO_normal_ and 1 for SO_early_. This approach was chosen for its simplicity, robustness, and ability to estimate probabilities of early senescence while providing interpretable coefficients. All models were run separately for beech and oak forests using data from all years.

#### 2.3.1 Univariate models

First, we fitted univariate models using either VPD_max_ or ψ_soil_ as predictors. As we assumed that SO_early_ occurs after a period of cumulative drought stress, we evaluated multiple averaging periods for both VPD_max_ and ψ_soil_. These periods extended from mid- to late August (the month with highest frequency of SO_early_) in biweekly time steps back to 1 April, the approximate start of the growing season. We selected the optimal averaging period for each predictor based on the highest F1-score (harmonic mean of precision and recall) from logistic regression models using 5-fold cross-validation with random splits. To address class imbalance resulting from a substantially lower number of occurrences of SO_early_ compared to SO_normal_, we applied weighting by inverse class frequencies during model fitting. Thresholds for drought stress triggering early senescence were derived from the logistic regression probability scores, with the 50% probability level defining the threshold.

#### 2.3.2 Bivariate models

We fitted bivariate models using both VPD_max_ and ψ_soil_ as predictors to assess the relative contributions of atmospheric and soil drought stress to SO_early_. Prior to modelling, predictors were standardized to zero mean and unit variance to allow direct comparison of effect sizes. The coefficients were derived across five random cross-validation splits and represent the change in the log-odds (log (p⁄(1 − p)), with p=probability of SO_early_) of SO_early_ per standard deviation increase in the predictor. Positive coefficients indicate that the likelihood of early senescence increases with the predictor, while absolute values reflect the contribution of each predictor to SO_early_.

We also explored bivariate models combining the most predictive averaging periods from summer and spring to evaluate the relative influence of drought at different seasonal stages. Interaction terms between predictors were tested for all bivariate models, but were retained only when they improved model performance (F1-score). To account for potential site-specific adaptation of trees to local conditions, we additionally tested models using standardized anomalies (z-scores) for both VPD_max_ and ψ_soil_ for the respective averaging periods, computed based on data for 2010–2023.

### 2.4 Early leaf senescence and mortality

To assess the relationship between early leaf senescence and subsequent canopy mortality (Q3), we used the cover fraction of standing deadwood in the tree canopy for Germany from 2018–2021 (Schiefer et al., 2025) derived from high-resolution UAV imagery combined with Sentinel-1 and Sentinel-2 data. Since our focus was on increases in canopy mortality following early leaf senescence, we calculated year-to-year differences in deadwood cover and retained only positive changes.

Negative changes could be related to both removal of dying trees or canopy regrowth and were thus neglected. We then aggregated the mortality increase maps to 1 km resolution by averaging across pixels within the respective forest types. We compared the distributions of mortality increase for forest pixels having experienced normal, early (for this analysis defined as SO between 15–31 August) or very early senescence (before 15 August) in the preceding year. Differences in median mortality increases between these groups were assessed using two-sided Mann–Whitney U tests. To evaluate the effect of repeated early senescence events on canopy mortality, we also compared forest pixels with no, single, or multiple occurrences of early leaf senescence in preceding years.

### 2.5 Drought sensitivity classification

Based on the previous analyses, we classified beech and oak forest pixels into different drought response types according to senescence timing, drought sensitivity, and mortality patterns (Q4). We therefore applied K-means clustering to eight variables: mean SO, SO variability, SO_early_ frequency, next-year mortality increase, and correlations of SO with VPD_max_ and ψ_soil_ in spring and summer.

These variables describe whether a forest pixel tends to senesce earlier or later, its interannual variability, responsiveness to atmospheric and soil drought stress, and vulnerability to drought-induced damage. The optimal number of clusters (k=3) was selected based on the inertia criterion (within-cluster sum of squares) while maintaining interpretability. To characterize and interpret the identified drought response types, we examined their associations with a range of stand, climatic, and edaphic variables. These included canopy cover, height, and height variability (as proxies for stand structure); timing of leaf maturation (date when VI reaches 80% of the annual amplitude in spring); seasonal and annual averages (calculated across all years) of temperature, climatic water balance (CWB), VPD, soil water potential at multiple depths (0–50, 0–100, 0–150 cm) and relative extractable water (REW) in the root zone; soil depth, texture (sand, silt, clay fractions), and bulk density; and topography (elevation, slope, exposition). All datasets were harmonized to 1 km resolution, and stand metrics were computed separately for beech- and oak-dominated forests. Data sources are listed in Table S2 (SI). To reduce redundancy and focus on the most informative variables, highly correlated environmental variables were grouped via hierarchical clustering, and only the variable with the highest mutual information with the drought response types was retained from each group.

## 3. Results

### 3.1 Senescence onset from satellite data

To evaluate the suitability of remotely sensed metrics for tracking senescence onset (SO), we first compared Sentinel-2–derived estimates against ground-based observations from ICP Forests Level II plots (Figure 1a-b). Remotely sensed SO as derived from different VIs and amplitude thresholds showed moderate agreement with field observations of coloration and litterfall onset (Table S3 and Table S4, SI). Among all VI–threshold combinations, the normalized difference red-edge index (NDRE) with the 80% seasonal amplitude threshold achieved the lowest, though still substantial RMSE, and moderate correlations for coloration (RMSE=18.1 days, Pearson’s r=0.41) and litterfall onset (RMSE=21.0 days, r=0.35). By contrast, NDVI showed particularly poor agreement with the field-recorded dates, with the best amplitude threshold for coloration being 85% (RMSE=31.9 days, r=0.20) and for litterfall being 80% (RMSE=31.2 days, r=0.26). A derivative-based estimation of SO (cf. Misra et al., 2016) did not improve results, nor did correlating subsequent observations of the phenology event (instead of the initial documented observation) with different remote sensing thresholds. Aggregating the remotely sensed SO estimates over larger areas generally improved the correlation with field observations, likely by reducing pixel-specific noise through spatial averaging (Figure 1c-d). However, the limited number of phenological observations in oak forests prevented identification of species-specific optimal VIs and thresholds. Consequently, SO maps derived from NDRE at the 80% threshold were used in all subsequent analyses.

**Figure 1:**
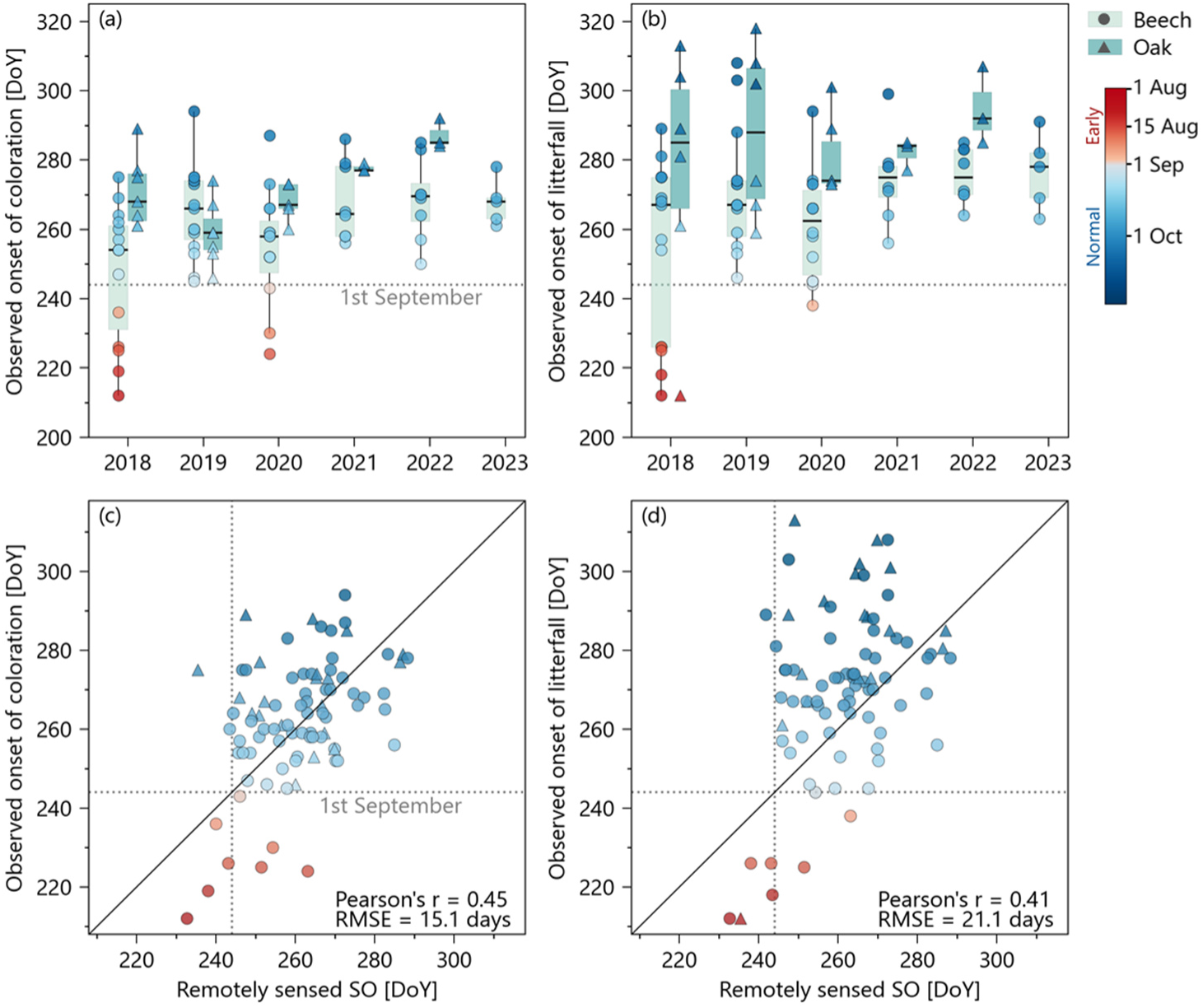
Observed onset of leaf senescence in beech and oak forests in Germany (2018-2023) derived from ground-based measurements and remote sensing. Ground data are from ICP Forests Level II plots (http://icp-forests.net) and include recordings of a) ‘infrequent or slight’ coloration (1-33% of the canopy affected), and b) ‘infrequent or slight’ litterfall. Note that the number of observed plots varied annually (n=5-16 for beech, n=3-7 for oak). Relationships with remotely-sensed senescence onset (SO) are shown as c) coloration observations vs. SO derived from Sentinel-2 Normalized Difference Red-Edge Index (NDRE) time series using a threshold of 80% of the seasonal amplitude and d) litterfall observations vs. the same NDRE-based SO. Note that the remotely sensed SO was averaged over an area of 40x40 km around the ICP Forests plots, therefore representing a regionalized senescence signal.

### 3.2 Patterns of early leaf senescence for 2018–2023 (Q1)

Leaf senescence onset in Germany’s beech- and oak-dominated forests showed strong spatial and temporal variability between 2018 and 2023. This resulted in marked differences in the prevalence of early leaf senescence (Figure 2). Early senescence was most widespread in 2018, affecting 42.9% of the forest area (13,817 km²), followed by 2019 (22.8%, 8,126 km²) and 2022 (20.1%, 7,195 km²). It was less prevalent in 2020 (13.7%, 5,010 km²) and 2021 (12.4%, 4,139 km²), and rare in 2023 (5.3%, 1,895 km²). Both beech and oak were affected in similar proportions each year, but the absolute area was consistently larger for beech forests, ranging from 8,769 km² in 2018 to 1,384 km² in 2023, compared with 5,048 km² and 512 km² for oak, respectively. Geographically, early senescence in 2018 was particularly widespread in beech forests of central Germany and in oak forests of the center and northeast. In 2019, early senescence hotspots shifted towards the northeast for both species, while in 2020, they clustered in the central-west. In subsequent years, patterns were more scattered across the country. The timing of early senescence also differed: an average of 47% of early-senescing beech and 65% of early-senescing oak forest began senescing before mid-August. Across the full study period, only 28.8% of the beech forests in Germany never showed early leaf senescence, while 47.3% experienced it once, 20.0% twice, and 3.5% three times; more frequent occurrences were rare (<1%) (Figure S1a, SI). For oak, 56.0% remained unaffected, 34.5% senesced early once, 7.7% twice, and 1.3% three times, while higher recurrence was very rare (<0.1%) (Figure S1b, SI).

**Figure 2:**
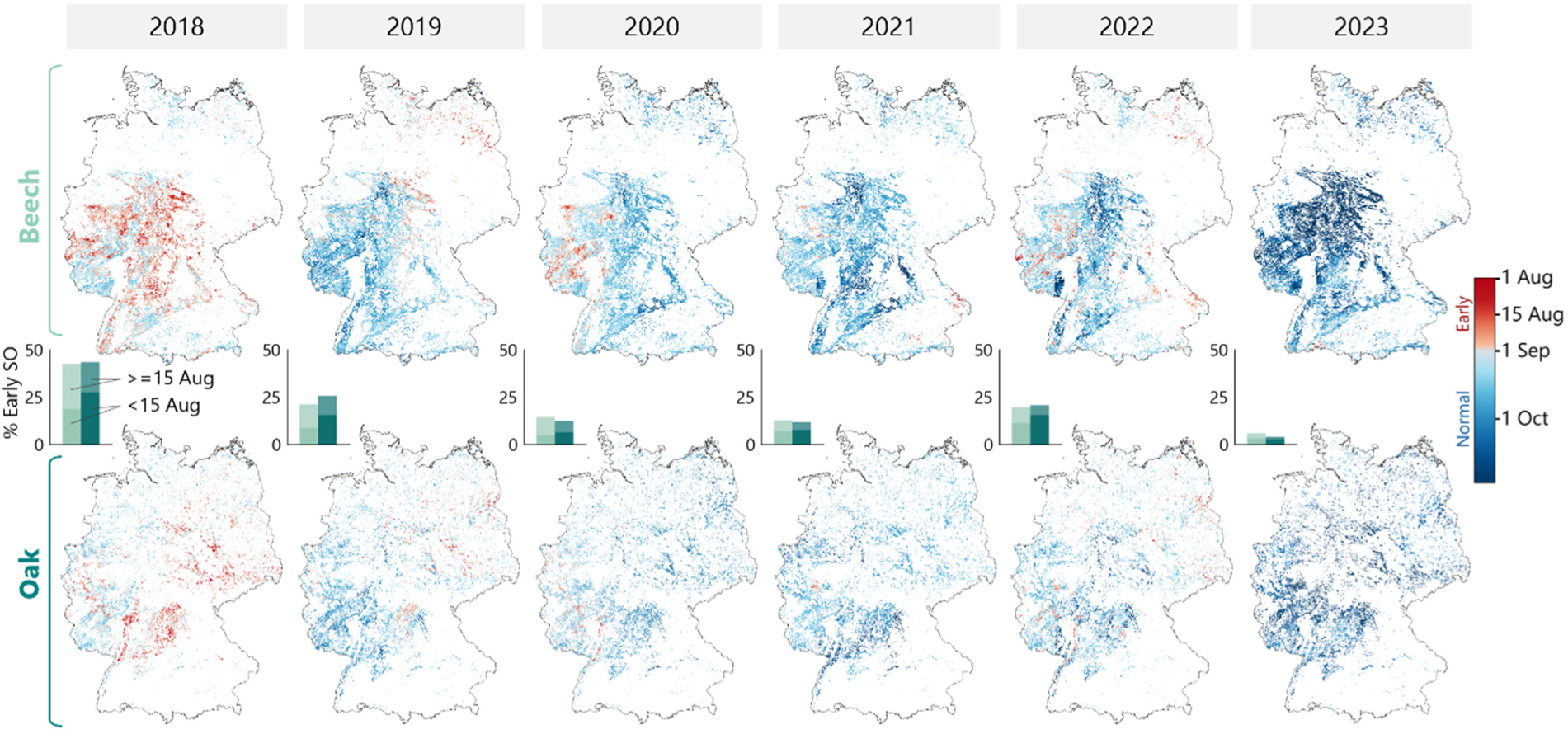
Maps showing the senescence onset (SO) for beech- and oak-dominated forests in Germany at 1 km resolution from 2018-2023. The colors represent the average SO of all 10 m pixels within each 1 km grid cell that are located in stands dominated by the respective species. Red colors indicate early senescence onset (SO_early_), defined as SO before DoY 244, i.e., 1 September in non-leap years. Bar plots show the percentage of SO_early_ within the beech (light green) and oak forest areas (dark green), respectively. Different shadings distinguish between areas with SO_early_ before and after 15 August.

### 3.3 Role of atmospheric and soil drought stress (Q2)

#### 3.3.1 Cumulative periods and thresholds of drought stress

We found that SO_early_ was related to atmospheric and soil drought indicators, but with different timing and duration of the most predictive period. For VPD_max_, the critical windows spanned six weeks from early July to mid-August (Figure 3a). Generally, periods covering these two summer months provided the best model performance, while shorter windows, such as only two weeks in August, performed worse (Figure 3a, and Table S5 in SI). The average VPD_max_ between 1 July and 15 August associated with a >50% probability of SO_early_ (logistic regression classification threshold) was 1.89 kPa in beech forests and 2.06 kPa in oak forests (Figure 3c).

**Figure 3:**
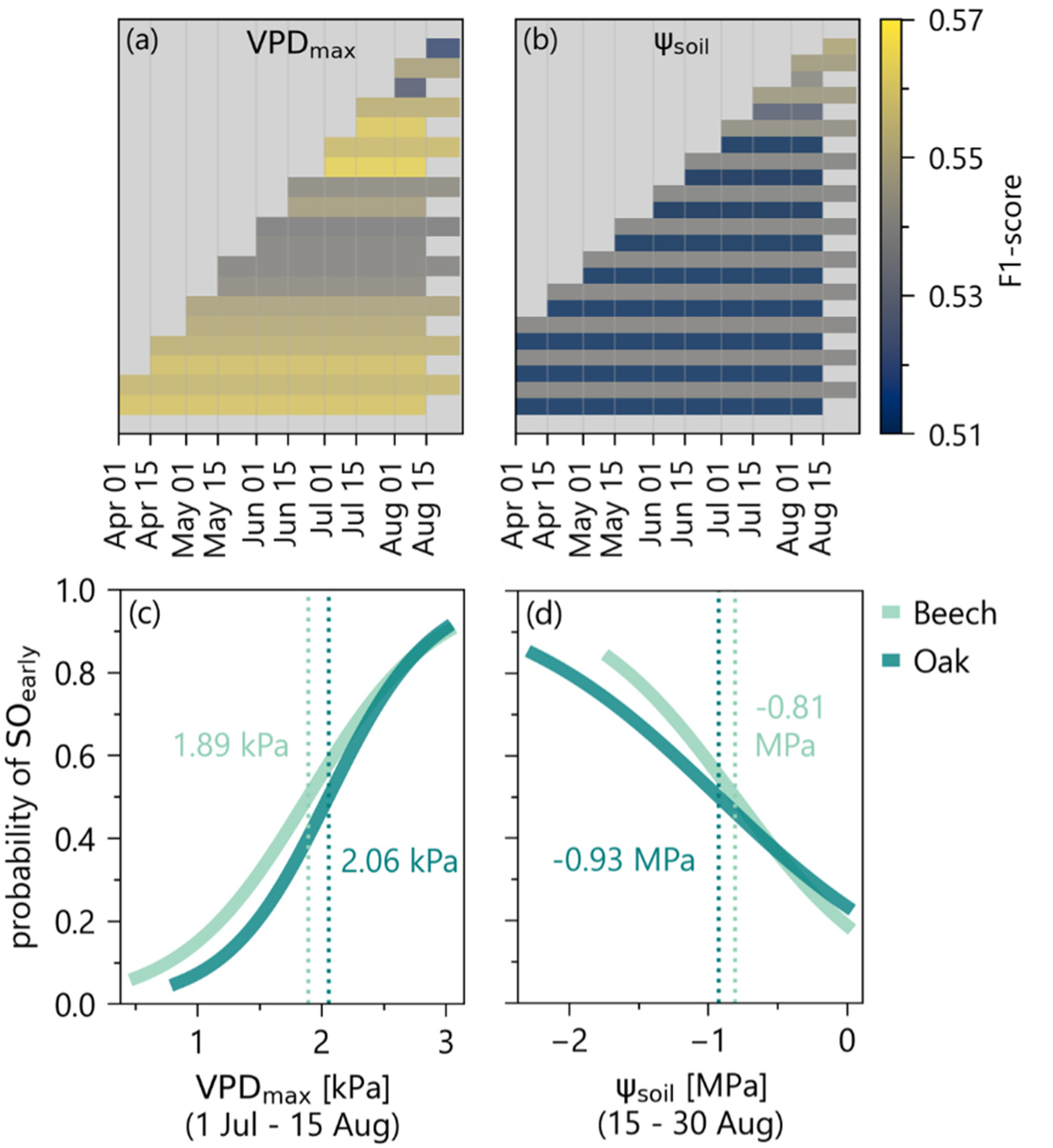
Comparison of cumulative periods of atmospheric and soil drought stress for predicting SO_early_, and modelled probability of SO_early_ for selected cumulative periods. a-b) Univariate logistic regression model performance (F1-score) across both species depending on different averaging time periods for VPD_max_ and ψ_soil_. c-d) The estimated probability of SO_early_ in beech and oak based on the most predictive averaging time periods. Vertical dotted lines show threshold values where the predicted probability of SO_early_ exceeds 50%.

For ψ_soil_, the two-week period from mid- to late August best captured the conditions triggering early senescence (Figure 3b). Time windows ending earlier in mid-August were less predictive, and extending the averaging period back to spring (1 April) also reduced model performance (Figure 3b, and Table S5 in SI). Over the second half of August, ψ_soil_ in early senescing stands (>50% probability of SO_early_) dropped below -0.81 MPa within the 1.4 m root zone in beech forests and below -0.93 MPa within the 1.8 m root zone in oak forests (Figure 3d).

Overall, the probability of early senescence was slightly higher in beech than in oak under comparable stress levels. Predictive performance was similar across indicators, with F1-scores of 0.56 (beech) and 0.57 (oak) for VPD_max_, and 0.57 (beech) and 0.54 (oak) for ψ_soil_.

#### 3.3.2 Relative importance and seasonal timing of soil versus atmospheric drought

We tested if combining VPD_max_ and ψ_soil_ within and across seasons improves the prediction of SO_early_ using bivariate logistic regression models.

Considering summer conditions only, we found that using VPD_max_ (1 Jul – 15 Aug; hereafter VPD_max,summer_) and ψ_soil_ (15–30 Aug; ψ_soil, summer_) together did not improve overall performance (F1-score=0.57 for both species), but the predictors differed in importance. Model coefficients suggest that ψ_soil, summer_ dominates in beech forests, and VPD_max,summer_ in oak forests (Figure 4a). Density plots support this, showing a more distinct separation in ψ_soil, summer_ values between early and normal senescing forests in beech compared to oak (Figure 4g), although oak forests generally experienced lower ψ_soil, summer_. When using standardized anomalies (see section 2.3.2), ψ_soil, summer_ gained importance for oak as well (Figure S2a, SI).

**Figure 4:**
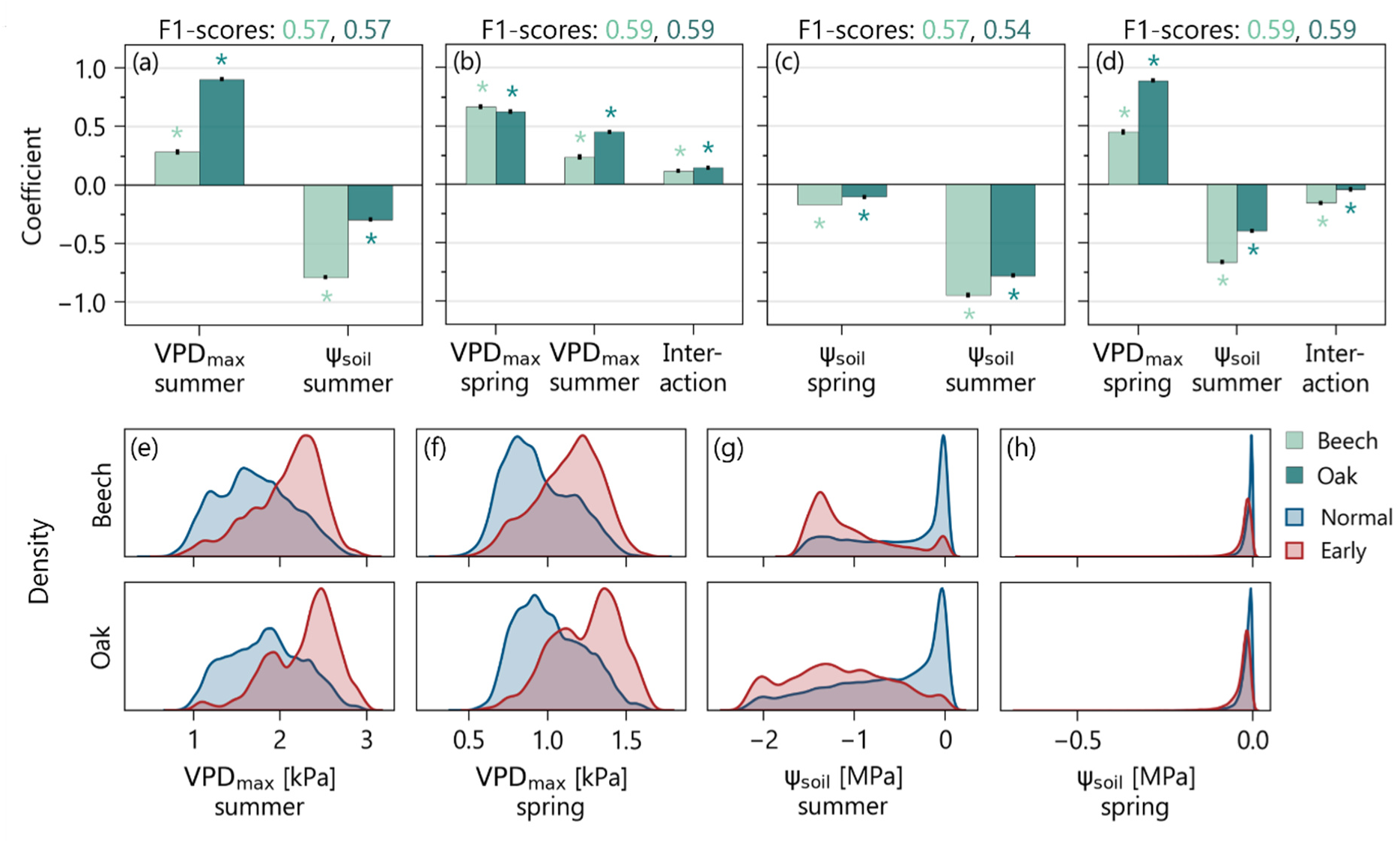
Different combinations of atmospheric and soil drought stress indicators were used to explain early leaf sensecence in bivariate models. a-d) Coefficients (effect sizes) of VPD_max,summer_ (1 Jul – 15 Aug), VPD_max,spring_ (15 Apr – 30 May), ψ_soil,summer_ (15–30 Aug)_,_ ψ_soil,spring_ (15–30 May) in four different models. Black error bars indicate the standard deviation of the coefficients across five cross-validation folds. Asterisks denote statistically significant coefficients. e-h) Probability density distributions of spring and summer VPD_max_ and ψ_soil_ for normal and early senescing forest pixels in beech (top row) and oak (bottom row).

Next, we combined spring and summer predictors. The most predictive spring windows were – similar to the summer periods – six weeks for VPD_max_ (15 Apr – 30 May; VPD_max,spring_), and two weeks for ψ_soil_ (15–30 May; ψ_soil, spring_) (Table S6, SI).

Combining VPD_max,summer_ (Figure 4e) and VPD_max,spring_ (Figure 4f) slightly raised the F1-score to 0.59 for both species. Coefficients indicated that elevated VPD in spring is particularly important (Figure 4b), with a positive interaction term suggesting that high atmospheric demand in both seasons compounds senescence risk. Models using standardized anomalies showed similar results (Figure S2b, SI).

In contrast, ψ_soil, spring_ contributed little to the prediction of SO_early_ when combined with ψ_soil, summer_ (Figure 4c) (F1-score=0.57 for beech, 0.54 for oak). Overlapping distributions of ψ_soil, spring_ for early and normal senescing forests confirm that soil water potential differences were minor at this time of the year (Figure 4h). Using anomalies instead of absolute values further emphasized the stronger role of ψ_soil, summer_ for both species (Figure S2c, SI).

Pairing VPD_max,spring_ with ψ_soil, summer_ yielded similar performance as the dual VPD_max_ model (F1-score=0.59 for both species). Here, ψ_soil, summer_ was more important for beech, whereas oak showed stronger effects of VPD_max,spring_ when using absolute values (Figure 4d), but had equal contributions of both predictors when using anomalies (Figure S2d, SI).

Lastly, we used spring and summer daily maximum temperatures instead of VPD_max_, which yielded the highest F1-scores (0.62 for beech, 0.61 for oak), indicating that direct heat stress may contribute to early senescence. However, it was not possible to separate the independent effects of VPD and temperature, as both variables were highly correlated (Pearson’s r=0.89 for daily values, 0.95 for seasonal averaging periods).

Overall, our results highlight that early senescence is primarily driven by the seasonal interplay of drought stress, with spring VPD amplifying the risk and summer soil water deficits acting as critical thresholds, albeit with species-specific sensitivities.

### 3.4 Mortality in subsequent years (Q3)

Early senescence was generally associated with increased canopy mortality in the next year (Figure 5a-c). Across both species and most years, the median increase in mortality was significantly larger (p<0.05) in early and very early senescing stands compared to normal ones, with the exception of very early senescing beech in 2019. In beech forests, early senescence (15–31 August) was followed by a 4.8–4.9 percentage points (pp) increase in mortality (expressed as percentage cover of the respective forest pixel), compared to 3.9–4.5 pp in normal senescing stands. In oak forests, early senescence was linked to a 4.3–6.6 pp mortality increase, compared to 4.0–5.5 pp after normal senescence. Very early senescence (before 15 August) amplified this effect, with mortality increases up to 5.8 pp in beech and 7.9 pp in oak (in 2019 with early senescence recorded in 2018).

**Figure 5:**
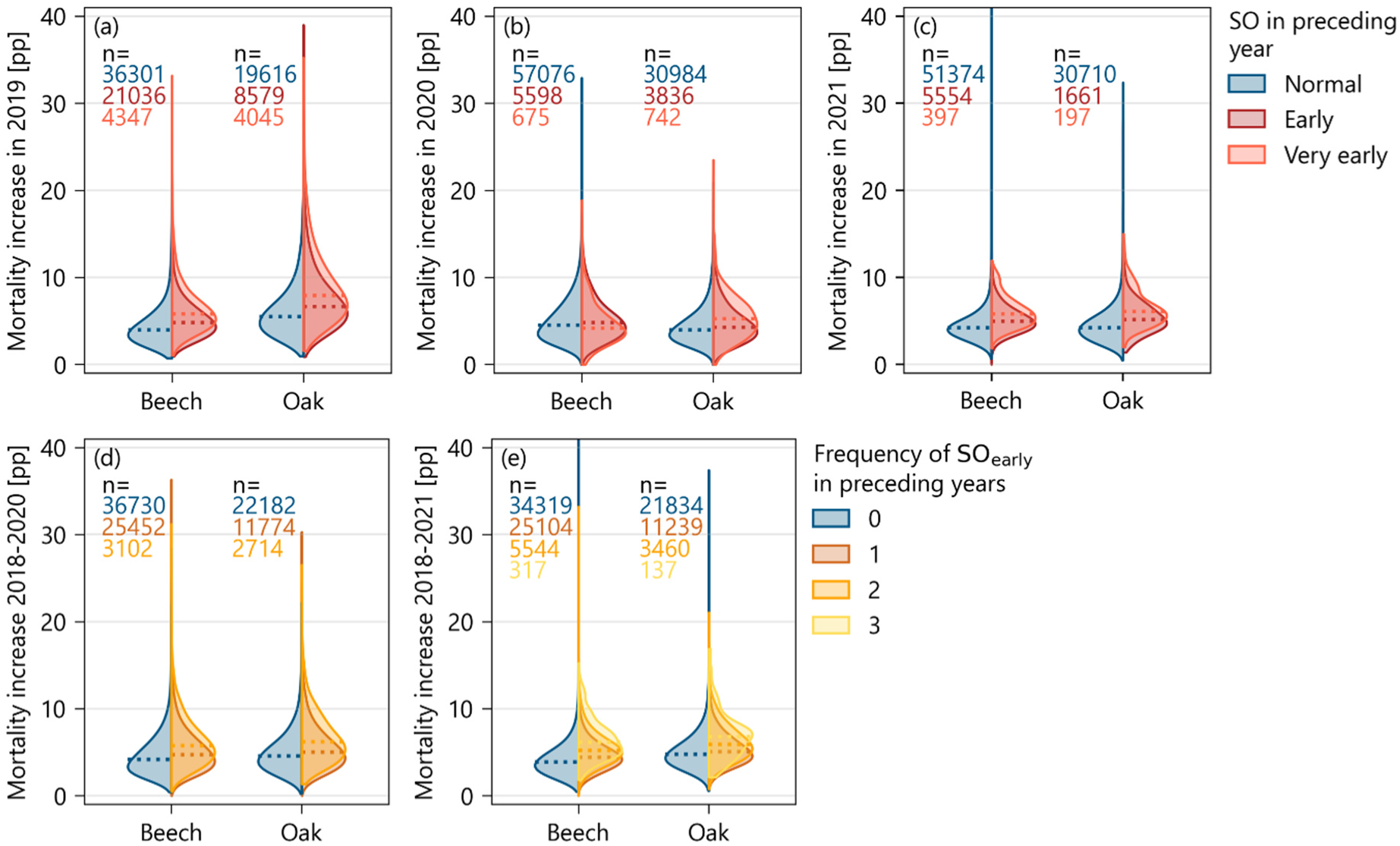
Mortality following early leaf senescence in beech and oak forests in Germany. a-c) Pixel-level increases in next-year canopy mortality cover in percentage points from Schiefer et al. (2023) for forest pixels with normal, early, or very early (SO before 15 August) senescence in the preceding year. Since mortality data only cover 2018-2021, the analysis is limited to the relationship between SO in 2018, 2019, and 2020 and mortality in the following year. Numbers denote sample sizes; horizontal dotted lines indicate medians of the distributions. Mortality effects beyond one year after early senescence are shown in Figure S3 (SI). d-e) Increases in canopy mortality in 2020 and 2021 (compared to 2018) after multiple early senescence events vs. no or only one occurrence in preceding years.

Not only the early timing of leaf senescence was linked to next-year tree mortality, but also the frequency of early senescence across several preceding years (Figure 5d-e). Forests with multiple early senescence events showed significantly larger increases in canopy mortality (p<0.05) than those with only a single occurrence, in both species and all investigated years. For example, in 2020, mortality relative to 2018 increased by 5.7 pp in beech and 6.2 pp in oak when early senescence occurred in both 2018 and 2019, compared to 4.7 pp and 5.0 pp, respectively, after only one occurrence. Three consecutive events further elevated mortality, increasing by 6.2 pp in beech and 6.8 pp in oak in 2021, compared to 4.4 pp and 5.0 pp in forests with only one event.

### 3.5 Forest drought response types (Q4)

The high spatial variability in the occurrence of early leaf senescence indicates that drought sensitivity differs markedly among forest sites. Using K-means clustering, we identified three distinct drought response types that were highly consistent across both species (Figure 6, Figure 7):

1. **Resistant** type – Forests with comparably low interannual variability in senescence onset (Figure 6b, 7b), rare early senescence (Figure 6c, 7c), and low mortality (Figure 6d, 7d). Their senescence timing showed little to no sensitivity to either spring or summer VPD_max_ and ψ_soil_ (as indicated by near-zero correlations with SO, Figure 6e-h, 7e-h).
2. **Sensitive** type – Forests with higher variability in senescence timing, but still low frequency of early senescence and low mortality. However, their senescence timing is highly sensitive to both spring and summer drought indicators, pointing to potential vulnerability under future drought conditions.
3. **Vulnerable** type – Forests with high variability in senescence onset, frequent early senescence, and elevated mortality. Their senescence timing is strongly influenced by VPD_max_ and ψ_soil_, particularly in summer, indicating both high sensitivity and exposure to drought stress.

**Figure 6:**
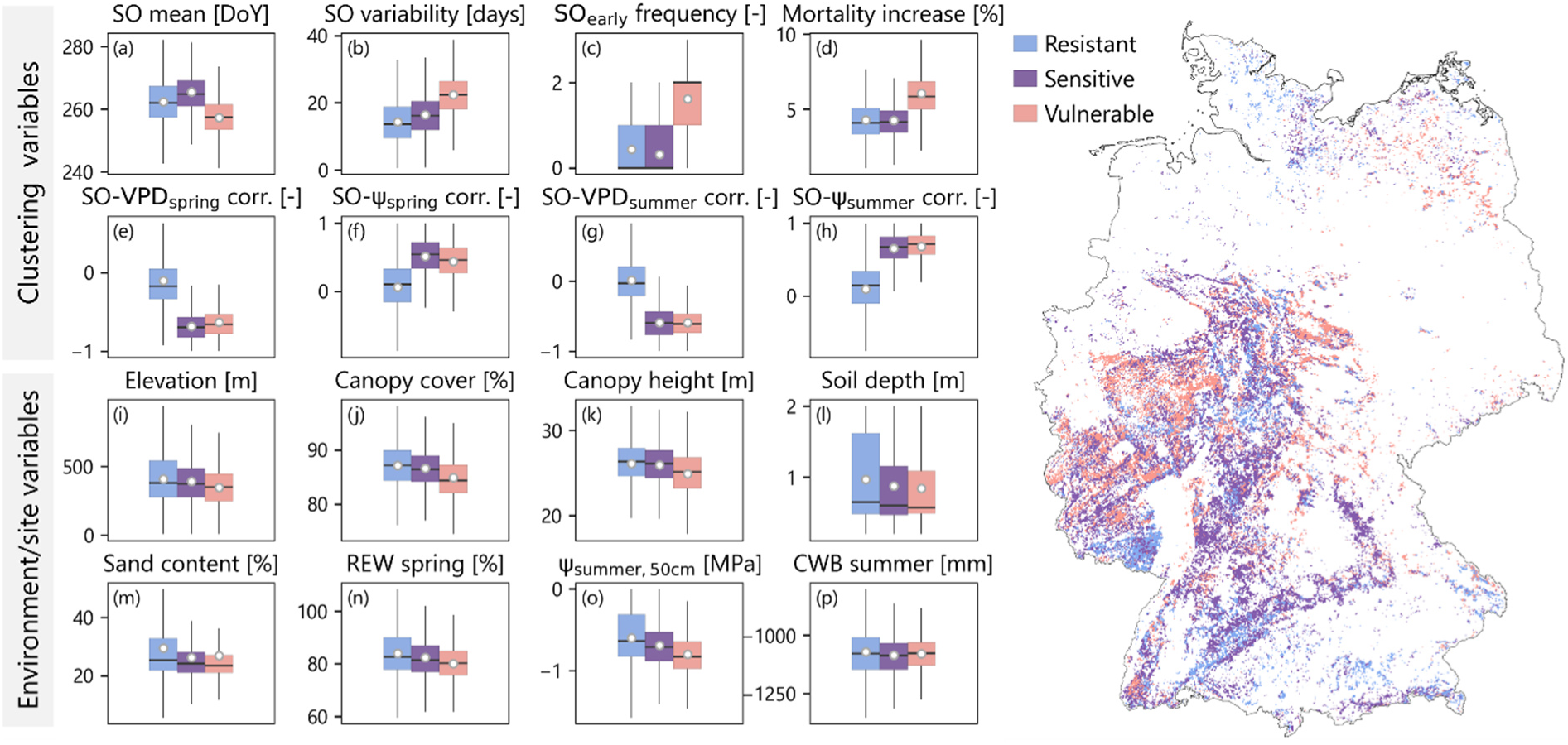
Clustering of beech forest pixels based on senescence timing, drought sensitivity (correlation between SO and VPD_max_ or ψ_soil_), and mortality patterns. Eight variables (a-h) were used to categorize the three groups: 1) Resistant (blue) – low drought sensitivity of SO timing (e-h) and SO_early_ frequency (c); 2) Sensitive (violet) – high SO–drought sensitivity but low SO_early_ frequency; 3) Vulnerable (pink) – high SO–drought sensitivity, high SO_early_ frequency, and increased mortality. Boxplots (i-p) illustrate how additional environmental variables differ across clusters. Displayed variables were selected based on hierarchical clustering and mutual information with the drought response types. The map depicts the spatial distribution of drought response types across Germany’s beech forests.

**Figure 7:**
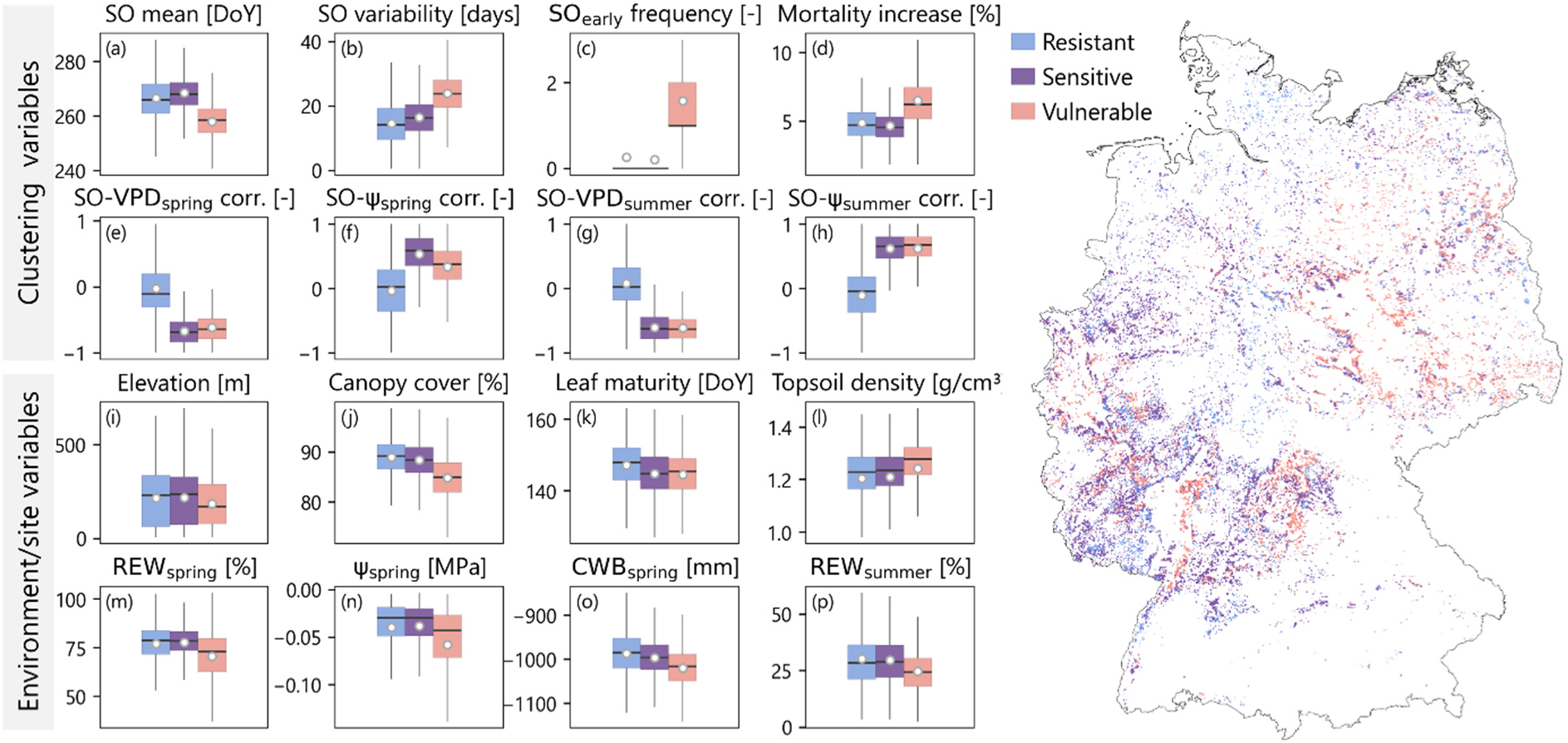
Same as Figure 6, but for oak forests. Note that the environmental variables (i-p) were selected based on hierarchical clustering and mutual information with the drought response types.

In beech forests, the resistant response type occurs mainly in the far north, along the lower mountain ranges and in the pre-Alpine region of southern Germany (Figure 6). These stands are generally located at medium high elevations, have closed canopies and tall trees, and often deep soils with higher sand content (Figure 6i-m). They benefit from greater soil water availability, as reflected by higher relative extractable water (REW) in spring and less negative ψ_soil_ in summer (Figure 6n-o). In contrast, the vulnerable response type is predominantly located in central Germany, particularly in lowland areas. These forests tend to have lower canopy cover and height, and are typically found on shallower soils. As a result, they experience poor water availability, especially in the upper soil layer during summer. The sensitive response type occupies an intermediate position, with structural and site characteristics between the two extremes. Under current conditions, however, these stands do not yet experience critically low soil moisture that would trigger widespread early senescence or elevated mortality.

In oak forests, the resistant response type is more frequently found in the northwest, where wetter spring weather conditions ensure a relatively high soil moisture at the beginning of the growing season, which also leads to more favorable water availability in summer (Figure 7m-p). Similar to resistant beech stands, these forests have closed canopies, and show late leaf maturation in spring (Figure 7j-k). The vulnerable response type is more common in central and northeastern Germany, typically at lower elevations with compacted soils and increased water deficits already in spring and throughout summer (Figure 7l-p). The canopies of these forests are more open and reach leaf maturity earlier in the year. Sensitive response types are again situated in intermediate environmental conditions between resistant and vulnerable types.

In summary, both beech and oak forests show similar drought response gradients, with resistant stands in wetter or higher-elevation areas, vulnerable stands in drier lowlands with shallower or compacted soils, and sensitive stands occupying intermediate conditions, highlighting the interplay of soil, canopy structure, and local climate in modulating drought vulnerability.

## 4. Discussion

### 4.1 Senescence onset from satellite data

Our analysis showed moderate agreement between satellite-derived SO and field observations of coloration and litterfall. Discrepancies between the two datasets are expected, given their respective sources of uncertainty. The accuracy of remotely sensed SO is influenced by noise in the raw data, arising from cloud contamination and atmospheric effects, as well as by the necessary filtering, interpolation, and smoothing steps, which contribute to uncertainties in phenology estimates.

Understory and admixed tree species may additionally alter the signal. Field observations are similarly subject to uncertainty due to irregular assessments at the ICP Forests plots (minimum biweekly; Raspe et al., 2020) and flexible definitions of phenological events, which can affect 1–33% of the canopy. SO is generally more difficult to estimate than spring leaf-out and remains underrepresented in phenology studies (Li et al., 2023). Considerable variability in SO has been documented both across estimates by different observers (e.g., 19 days; Klosterman et al., 2014), and within populations of the same tree species (e.g., 19–33 days in oak, 26–28 days in beech; Delpierre et al., 2017). Due to the large within-canopy heterogeneity, remote sensing signals, even at 10 m resolution, primarily represent an averaged signal rather than individual tree variation.

Accurate calibration therefore requires regular, objective, and species-specific reference data – for example, a coloration percentage identified from UAV imagery across the tree crowns in a defined area (Krause & Sanders, 2024), or a percentage of the seasonal total leaf area from weekly littertrap collections. Despite these uncertainties, our results indicate that red-edge–based VIs such as NDRE are better suited to capture SO than the commonly used NDVI. It should be noted, however, that remote sensing alone cannot readily distinguish drought-induced from normal autumn senescence without incorporating timing information or additional drought stress metrics.

### 4.2 Patterns of early leaf senescence (Q1)

Our satellite-derived SO maps revealed that early leaf senescence affected both beech- and oak-dominated forests across multiple years. The most extensive occurrence in 2018 aligns with frequent reports of drought-related symptoms such as leaf curling, premature coloration and shedding, or even green leaf abscission by forestry institutions from several federal states (FVA-BW, 2018; LM, 2019a; NW-FVA, 2018; SBS, 2018). Reports also document strong fructification in both species, a resource-intensive process linked to elevated crown defoliation in beech (Eichhorn et al., 2005).

Intense masting may have contributed to the remotely sensed senescence signal by enhancing the apparent canopy browning. In some regions, extreme drought even triggered green fruit abortion, reflecting resource shortages and exacerbated physiological stress (LM, 2019a; Nussbaumer et al., 2020).

In subsequent years, our maps captured a reduced extent of early senescence and a geographic shift in affected areas. Forestry reports for 2019 describe deteriorating crown condition, with smaller beech leaves, early coloration in beech and marginal leaf browning in oak over the summer, and rising crown dieback (FVA-BW, 2019; LFE, 2019; LM, 2019b; NW-FVA, 2019). Regionally, drought-weakened trees experienced rising insect and pathogen activity, including bark necrosis and oak defoliators (FVA-BW, 2019; LFE, 2019; Thierfelder, 2020), indicating the emergence of compound stress interactions. This trend continued into 2020, with increased attacks by beech and oak bark beetles, early senescence on extreme sites, but locally also improved oak crown condition (FVA-BW, 2020; NW-FVA, 2020; Wald und Holz NRW, 2020). Detections of early senescence in 2019–2020 may thus reflect both immediate drought responses and progressive decline in chronically stressed stands. Lower prevalence of early senescence in 2021 aligns with wetter and cooler conditions, observations of later leaf unfolding, partial stabilization of crown condition, and rare reports of early senescence (FVA-BW, 2021; NW-FVA, 2021; Wald und Holz NRW, 2021). The widely dry and hot summer in 2022 was again reflected in more widespread satellite detections of early senescence, consistent with more frequent field observations, reported crown dieback in beech and increased pest-related damage in oak (FVA-BW, 2022; LFE, 2022; LM, 2022). In 2023, despite continued high levels of crown damage, our data showed very limited early senescence, which agrees with reports of favorable water supply reducing acute drought stress symptoms, while oak showed both partial crown regeneration and insect-related decline (FVA-BW, 2023; LM, 2023; NW-FVA, 2023). Overall, the satellite patterns correspond closely with ground-based, although not spatially explicit observations of drought stress symptoms, regionally amplified by fructification and biotic damage, highlighting that early senescence reflects both immediate physiological stress and longer-term vitality decline in Germany’s beech and oak forests.

### 4.3 Seasonal roles of soil and atmospheric drought (Q2)

Our analyses indicate that early leaf senescence was not triggered by instantaneous VPD but by sustained periods of high atmospheric demand that progressively depleted soil water reserves. The identified time windows — six weeks for VPD_max_ and the subsequent two weeks for ψ_soil_ — capture complementary stages of drought stress, and can to some extent be used interchangeably to explain early senescence occurrence. However, their relative predictive power differed between species, reflecting differences in the sensitivity to soil and atmospheric drought.

Beech was particularly sensitive to soil drought, consistent with its shallower rooting system and higher vulnerability to embolism formation (Jonard et al., 2011; Leuzinger et al., 2005), the likely cause for drought-induced defoliation (Walthert et al., 2021). Our model suggests that the probability of early senescence in beech exceeds 50% when ψ_soil_ in the root zone drops below -0.81 MPa. Despite numerous assumptions underlying the simulated soil water potential used in this study, this value closely matches the -0.8 MPa threshold previously identified by Walthert et al. (2021) for the onset of beech defoliation in field studies. Oak exhibited a lower threshold of -0.93 MPa, despite a deeper assumed root zone, reflecting the ability to sustain water transport under more stressful conditions. Indications for stronger senescence signals in oak under atmospheric drought may thus also reflect indirect effects, such as enhanced likelihood of insect outbreaks under hot, dry conditions (Haynes et al., 2022; Karolewski et al., 2007).

Spring atmospheric dryness further contributed to early senescence, consistent with legacy effects reported in previous studies (Bigler & Vitasse, 2021; Fu et al., 2014; Zuccarini et al., 2023). This has been related to carryover effects of an earlier spring phenology on leaf senescence, but different mechanisms have been proposed: Earlier leaf unfolding may accelerate soil water depletion, increasing the risk of critical soil water levels being reached earlier than usual during summer (Bigler & Vitasse, 2021). Constraints on leaf longevity after an early budburst and cumulative stress damage may shorten the functional leaf lifespan after a dry spring (Mariën et al., 2021; Zuccarini et al., 2023), while sink limitation following high early-season productivity may trigger earlier senescence via carbon storage feedbacks (Fu et al., 2014; Zani et al., 2020). Still, the key process through which spring conditions impact leaf senescence needs to be elucidated. We further hypothesize that dry springs may increase drought vulnerability later in the season by limiting the formation of new sapwood needed to sustain transpiration of the developed leaves (Bose et al., 2021; Etzold et al., 2022). Particularly in 2019 and 2020, drought stress occurred early in the growing season in many areas, resulting in strong radial growth reductions particularly in beech (Scharnweber et al., 2020; Thom et al., 2023), which may have constrained water transport and contributed to early leaf senescence in these years.

Our findings underline that combined spring and summer droughts, events expected to become more frequent across Europe (Spinoni et al., 2018), increase the likelihood of early leaf senescence. The thresholds identified here provide a basis for integrating early senescence into ecosystem models to better predict drought impacts on carbon fluxes and tree mortality. Recent advances in representing soil and plant hydraulics in such models (Nadal-Sala et al., 2024; Scheiter et al., 2024) now allow simulation of drought-induced tissue damage and leaf shedding, which could be informed by our results. While our analyses emphasize drought as the primary driver of early senescence, the strong correlation between VPD and temperature in our dataset prevented us from disentangling their independent effects, which should be addressed in future experimental work. Finally, we note that our models are strongly influenced by the exceptional drought of 2018, when early senescence was most widespread. Thus, the identified time windows indicate average stress duration and intensity needed to trigger senescence, rather than precise seasonal timings.

### 4.4 Mortality in subsequent years (Q3)

Our results indicate that both exceptionally early senescence and repeated early senescence across years are linked to increased canopy mortality, supporting the view that early senescence can serve as indicator of drought-induced vitality decline (Frei et al., 2022). Although the satellite-based mortality estimates used in this study may potentially not fully separate mortality from defoliation, the observed patterns align with physiological evidence: early senescing beech trees often show persisting embolisms and reduced secondary growth, limiting their ability to restore hydraulic function and predisposing them to canopy dieback in subsequent years (Arend et al., 2022; Braun et al., 2021; Neycken et al., 2024; Schuldt et al., 2020). In oak, persisting xylem embolisms are less likely to cause mortality as the ring-porous species forms new xylem annually, but repeated early senescence and drought might lead to a depletion of carbon reserves that can ultimately impair hydraulic function (Plotkin et al., 2021). Elevated oak mortality has been reported when drought coincides with repeated insect defoliation, emphasizing the importance of drought–insect interactions (Tanzer et al., 2025). The overall increases in canopy mortality observed in our study were modest, which might indicate that early leaf senescence does not always signal severe damage and may, in some cases, represent an acclimation strategy that allows short-term survival. Whether early senescence ultimately leads to whole-tree mortality or recovery likely depends on the severity and recurrence of defoliation, individual tree repair capacity, and site-specific growing conditions.

### 4.5 Drought sensitivity across German forests (Q4)

Our study revealed strong variability in leaf senescence responses to drought and associated mortality across beech- and oak-dominated forests in Germany, largely driven by local soil water availability. This pattern is consistent with global analyses showing greater drought sensitivity of leaf senescence in more arid regions (Li et al., 2025) and with local studies linking higher growth sensitivity to climate variability and more severe crown damage to soils with lower water storage capacity (Klesse et al., 2022). We found reduced drought sensitivity at higher elevations, where temperature extremes are likely buffered by sufficient precipitation that reduces exposure to drought, and previous studies also report delayed senescence under such conditions (Bigler & Vitasse, 2021; Lukasová et al., 2020). Conversely, our study emphasizes the increased drought vulnerability of forests in lowland areas, on shallower and drier soils with less favorable growing conditions that are reflected in lower canopy heights (cf. Klesse et al., 2022).

Additionally, our results revealed a higher drought susceptibility of beech on sites with low sand content – similarly, oak was more vulnerable on soils with increased bulk density. This corresponds to reports of increased drought damage on loamy, compacted soils that become cemented during prolonged drought, which can impair the fine root system (LFE, 2019; SBS, 2018). We further detected greater drought sensitivity of leaf senescence in less dense stands, which may seem counterintuitive, as lower competition should improve water availability. Yet, repeated observations of early leaf coloration along forest edges and in single exposed trees (FVA-BW, 2018; LFE, 2019; Wohlgemuth et al., 2020), indicate that higher irradiation and evaporative demand in open canopies can accelerate leaf aging and dehydration. We did not detect increased vulnerability on south-exposed slopes, likely due to the coarse resolution of the topographic data. Nevertheless, topography is expected to influence drought impacts not only via radiation but also through water redistribution effects.

In summary, our findings highlight that soil water availability is the primary determinant of drought vulnerability, but interactions with stand structure, topography, and other soil properties further modulate local responses. While factors like groundwater access and resource competition likely shape individual tree water status, these remain challenging to capture at large scales. Overall, our results showed that large areas of Germany’s beech- and oak-dominated forests are vulnerable or sensitive to drought, underscoring their critical exposure to future climate extremes.

## 5. Conclusions

Satellite-based mapping of early leaf senescence reveals spatially and temporally variable drought impacts in German beech- and oak-dominated forests. We show that species-specific sensitivities to sustained atmospheric and soil drought, combined with seasonal legacy effects, link cumulative drought stress to senescence response. Very early and recurrent early leaf senescence increase canopy dieback, establishing them as physiological early-warning signals of drought-induced decline and enabling targeted monitoring. Further, local soil conditions and stand structure mediate drought vulnerability, highlighting sensitive regions as a focus for adaptive forest management. By identifying species-specific critical thresholds, our study supports incorporating drought-driven senescence dynamics into ecosystem models, which will likely improve projections of carbon cycling and forest mortality under future climate extremes. Overall, our findings underscore the need to anticipate and manage increasing drought vulnerability in temperate forests of Central Europe.

## Acknowledgements

We thank Paul Schmidt-Walther (DWD) for kindly providing soil water potential and relative extractable water data from LWF-Brook90. Evaluation of satellite-derived senescence onset was based on data collected by partners of the official UNECE ICP Forests Network (http://icp-forests.net/contributors). Part of the data was co-financed by the European Commission (Data accessed on 22/01/2025). We are grateful to Tanja Sanders for her assistance with the acquisition of the ICP Forests data and her helpful comments on related aspects of the manuscript. Satellite data processing was performed on the EO-Lab platform, operated by DLR and funded by the German Federal Ministry for Economic Affairs and Climate Action (BMWK). We used ChatGPT-5 (OpenAI) to support coding and improve readability and flow of the manuscript draft. This study was funded through the Helmholtz Initiative and Networking fund (W2/W3-156).

## Notes

### Competing Interest Statement

The authors have declared no competing interest.

