## supplemental file for "Soil and atmospheric drought trigger early leaf senescence and increase subsequent canopy mortality risk in temperate forests"

*Table S1: Vegetation indices (VIs) evaluated for estimating senescence onset. Wavelength regions correspond to Sentinel-2 bands as follows: Red: B4, Rededge1: B5, Rededge2: B6, Rededge3: B7, NIR: B8.*

| VI | Full name | Formula | Reference |
|---|---|---|---|
| NDRE | Normalized Difference Red-Edge Index | $\frac{Rededge2 - Rededge1}{Rededge2 + Rededge1}$ | (A. Gitelson & Merzlyak, 1994) |
| NDVIre1 | Normalized Difference Vegetation Index red-edge | $\frac{NIR - Rededge1}{NIR + Rededge1}$ | (A. Gitelson & Merzlyak, 1994) |
| CRE | Chlorophyll red-edge index | $\frac{Rededge3}{Rededge1} - 1$ | (A. A. Gitelson et al., 2003) |
| NDVI | Normalized Difference Vegetation Index | $\frac{NIR - Red}{NIR + Red}$ | (Tucker, 1979) |

*Table S2: Environmental and site variables used in the analysis with their respective sources.*

| Variable | Spatial resolution | Temporal resolution | Source |
|---|---|---|---|
| Canopy cover | 10 m | Single year | Copernicus Tree Cover Density 2018 (Copernicus, 2024) |
| Canopy height | 10 m | Single year | ETH Global Canopy Height 2020 (Lang et al., 2022) |
| Temperature | 1 km | Hourly | HOSTRADA – Hourly grids of high-resolution variables for Germany (DWD, 2024a) |
| CWB | 1 km | Monthly | Calculated from monthly sums of evapotranspiration and precipitation (DWD, 2024b) |
| VPD | 1 km | Hourly | Calculated from HOSTRADA temperature and dew point temperature (DWD, 2024a, 2024b) |
| Soil water potential | 1 km | 8-daily | Soil water potential in the root zone under beech and oak (P. Schmidt-Walter, personal communication, April 2025) calculated with LWF-Brook90 (Schmidt-Walter et al., 2020) |
| Relative extractable water | 1 km | 8-daily | Relative extractable water in the root zone under beech and oak (P. Schmidt-Walter, personal communication, April 2025) calculated with LWF-Brook90 (Schmidt-Walter et al., 2020) |
| Soil texture | 250 m | - | SoilGrids250m (Poggio et al., 2021) |
| Soil bulk density | 250 m | - | SoilGrids250m (Poggio et al., 2021) |
| Soil depth | 250 m | - | Soil depth in Germany / Physiologische Gründigkeit der Böden Deutschlands (BGR, 2015) |
| Topography | 1 km | - | ASTGDEM v3 digital elevation model (Sismanidis, 2024) |

*Table S3: Pearson's r and RMSE between ground-observations of **leaf coloration** onset and remotely sensed senescence onset for different vegetation indices (VIs) and different seasonal amplitude thresholds.*

| | **90%** | | **85%** | | **80%** | | **75%** | | **70%** | |
|---|---|---|---|---|---|---|---|---|---|---|
| **VI** | **r** | **RMSE** | **r** | **RMSE** | **r** | **RMSE** | **r** | **RMSE** | **r** | **RMSE** |
| NDRE | 0.19 | 35.4 | 0.27 | 23.5 | **0.41** | **18.3** | 0.40 | 22.2 | 0.38 | 27.1 |
| NDVIre1 | 0.21 | 30.0 | 0.27 | 22.4 | 0.41 | 19.4 | 0.41 | 23.2 | 0.38 | 28.2 |
| CRE | 0.16 | 52.0 | 0.18 | 41.5 | 0.17 | 33.8 | 0.25 | 25.5 | 0.31 | 20.2 |
| NDVI | 0.11 | 33.0 | 0.20 | 31.9 | 0.25 | 33.2 | 0.34 | 36.8 | 0.34 | 41.2 |

*Table S4: Pearson's r and RMSE between ground-observations of **litterfall** onset and remotely sensed senescence onset for different vegetation indices (VIs) and different seasonal amplitude thresholds.*

| | **90%** | | **85%** | | **80%** | | **75%** | | **70%** | |
|---|---|---|---|---|---|---|---|---|---|---|
| **VI** | **r** | **RMSE** | **r** | **RMSE** | **r** | **RMSE** | **r** | **RMSE** | **r** | **RMSE** |
| NDRE | 0.12 | 41.4 | 0.22 | 28.3 | **0.35** | **21.0** | 0.36 | 22.4 | 0.38 | 25.1 |
| NDVIre1 | 0.19 | 35.3 | 0.22 | 26.6 | 0.35 | 21.5 | 0.36 | 23.0 | 0.37 | 26.1 |
| CRE | 0.15 | 58.0 | 0.16 | 47.4 | 0.14 | 39.5 | 0.24 | 30.3 | 0.26 | 24.5 |

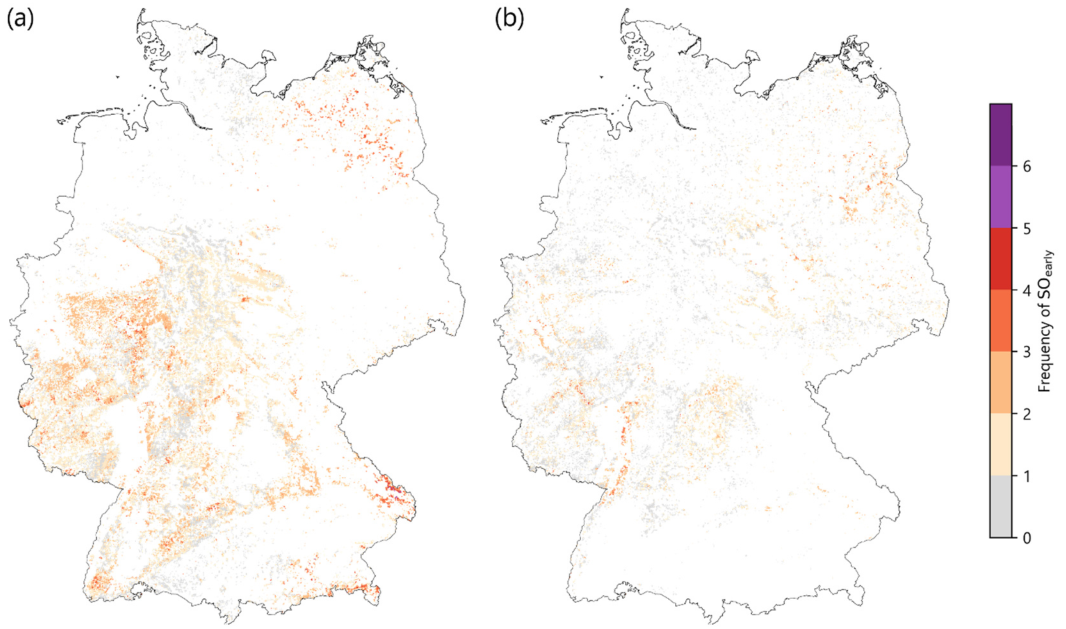


*Figure S1: Frequency of early leaf senescence in (a) beech and (b) oak forests between 2018 and 2023.*

*Table S5: Univariate logistic regression model performance (F1-score) depending on* ***different averaging time periods ending in August*** *for $VPD_{max}$ and $\psi_{soil}$. across beech and oak forests. Numbers indicate the start and end month of the averaging periods.*

| **$VPD_{max}$** | | **$\psi_{soil}$** | |
|---|---|---|---|
| **period** | **F1-score** | **period** | **F1-score** |
| 8_mid-8_end | 0.52 | 6_mid-8_mid | 0.51 |
| 8_start-8_mid | 0.53 | 6_start-8_mid | 0.51 |
| 6_start-8_end | 0.54 | 7_start-8_mid | 0.51 |
| 6_start-8_mid | 0.54 | 5_mid-8_mid | 0.51 |
| 5_mid-8_end | 0.54 | 5_start-8_mid | 0.51 |
| 5_mid-8_mid | 0.54 | 4_mid-8_mid | 0.51 |
| 6_mid-8_end | 0.54 | 4_start-8_mid | 0.51 |
| 6_mid-8_mid | 0.55 | 7_mid-8_mid | 0.53 |
| 8_start-8_end | 0.55 | 6_start-8_end | 0.54 |
| 5_start-8_end | 0.55 | 6_mid-8_end | 0.54 |
| 5_start-8_mid | 0.55 | 5_mid-8_end | 0.54 |
| 4_mid-8_end | 0.56 | 5_start-8_end | 0.54 |
| 7_mid-8_end | 0.56 | 4_start-8_end | 0.54 |
| 4_start-8_end | 0.56 | 4_mid-8_end | 0.54 |
| 7_start-8_end | 0.56 | 8_start-8_mid | 0.54 |
| 4_mid-8_mid | 0.56 | 7_start-8_end | 0.55 |
| 4_start-8_mid | 0.56 | 7_mid-8_end | 0.55 |
| 7_mid-8_mid | 0.56 | 8_start-8_end | 0.55 |
| 7_start-8_mid | 0.57 | 8_mid-8_end | 0.55 |

*Table S6: Univariate logistic regression model performance (F1-score) depending on* ***different averaging time periods in spring*** *for $VPD_{max}$ and $\psi_{soil}$. across beech and oak forests. Numbers indicate the start and end month of the averaging periods.*

| **$VPD_{max}$** | | **$\psi_{soil}$** | |
|---|---|---|---|
| **period** | **F1-score** | **period** | **F1-score** |
| 5_mid-6_end | 0.44 | 4_mid-4_end | 0.48 |
| 5_start-6_end | 0.48 | 4_start-4_end | 0.48 |
| 5_mid-6_mid | 0.49 | 4_mid-5_mid | 0.50 |
| 5_mid-5 end | 0.51 | 4_start-5_mid | 0.50 |
| 4_mid-4_end | 0.52 | 5_mid-6_end | 0.51 |
| 5_start-6_mid | 0.52 | 5_start-6_end | 0.51 |
| 5_start-5_end | 0.55 | 4_mid-5_end | 0.53 |
| 4_start-4_end | 0.55 | 4_start-5_end | 0.53 |
| 4_start-5_end | 0.57 | 5_start-5_end | 0.54 |
| 4_start-5_mid | 0.57 | 5_start-6_mid | 0.55 |
| 4_mid-5_mid | 0.58 | 5_mid-6_mid | 0.55 |
| 4_mid-5_end | 0.58 | 5_mid-5_end | 0.55 |

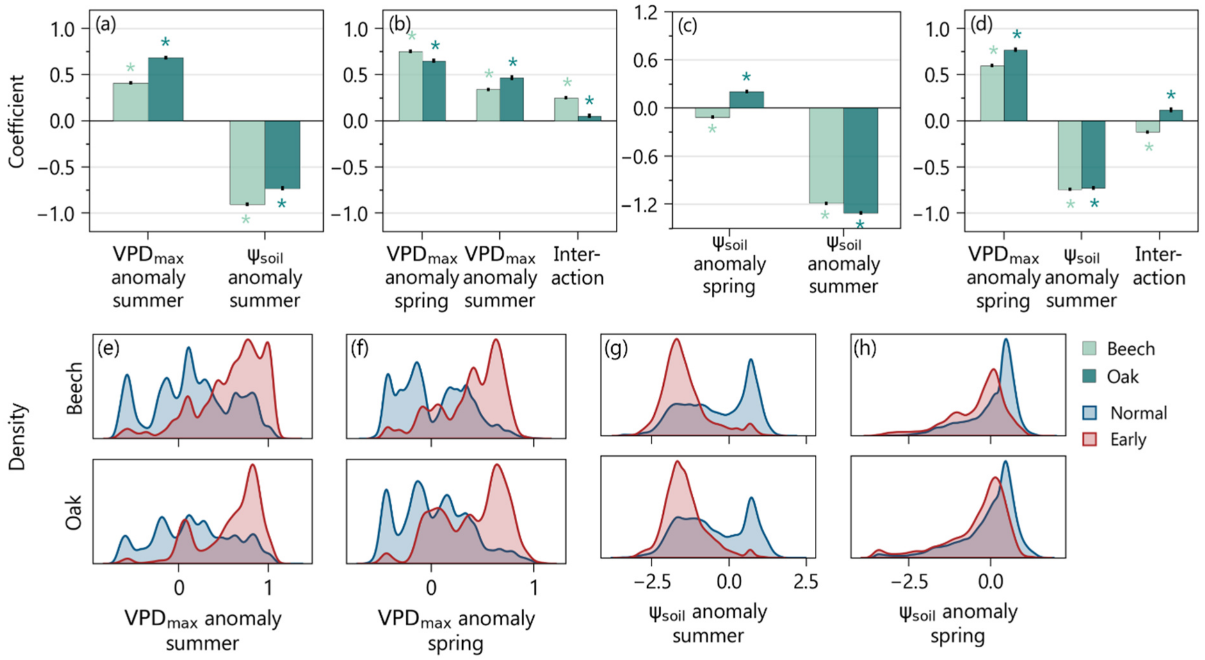


*Figure S2: Different combinations of the standardized anomalies of atmospheric and soil drought stress indicators were used to explain early leaf senescence in bivariate models. a-d) Coefficients (effect sizes) of $VPD_{max,summer}$ (1 Jul – 15 Aug) anomaly, $VPD_{max,spring}$ (15 Apr – 30 May) anomaly, $\psi_{soil,summer}$ (15–30 Aug) anomaly, $\psi_{soil,spring}$ (15 –30 May) anomaly in four different models. Black error bars indicate the standard deviation of the coefficients across five cross-validation folds. Asterisks denote statistically significant coefficients. e-h) Probability density distributions of spring and summer $VPD_{max}$ and $\psi_{soil}$ anomalies for normal and early senescing forest pixels in beech (top row) and oak (bottom row).*

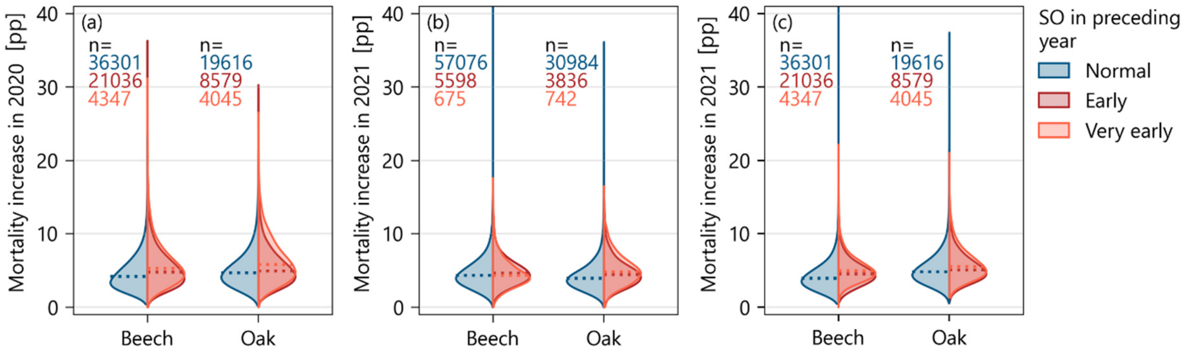


*Figure S3: Canopy mortality increase (difference in fractional cover expressed in percentage points, pp) two years and three years after early leaf senescence in beech and oak forests in Germany. a) shows mortality increases in 2020 after early senescence in 2018, b) shows mortality increases in 2021 after early senescence in 2019 and c) shows mortality increases in 2021 after early senescence in 2018.*
